# Development of a Multidose Irradiation Protocol for Clonogenic Assays in Cell Culture Plates

**DOI:** 10.64898/2026.06.05.730440

**Authors:** Bernarda Huilén Medina, Pablo Andrés, Lara Negrin, Luisa V. Biolatti, Sara Destri, Laura Mazzitelli-Fuentes

## Abstract

**Purpose:** *In vitro* experimental radiobiology is a fundamental tool for understanding the cellular and molecular mechanisms involved in the response to ionizing radiation. Conventional experimental designs require separate irradiations to achieve different absorbed doses, thereby introducing experimental variability between irradiation sessions due to inter-session variability in both culture conditions and irradiation geometry. Here, we developed a multidose irradiation system for multiwell cell culture plates that enables the simultaneous delivery of three distinct dose levels within a single plate, thereby reducing resource consumption and operating time

**Methods:** The system was designed using a clinical linear accelerator delivering 6 MV X-rays and a 3D conformal irradiation approach based on CT imaging. Dose calculations for 200, 400 and 600 cGy were performed using Monaco Version 5.11. The irradiation geometry was optimized to achieve distinct and well-separated dose regions while preserving dose uniformity within each dose level. Treatment planning showed good agreement between estimated and prescribed dose levels, with mean dose deviation ranging from 0.35-1.63%. Physical verification using TLDs and radiochromic films demonstrated high dosimetric accuracy, showing deviations of 0.5–1.0% and 0.57–3.0%, respectively, expressed as the deviation of the measured mean dose from the nominal administered dose levels. Biological assessment included clonogenic assays in human tumor cell lines and a metabolic assay. Clonogenic assays showed a high concordance between the multidose and single-dose irradiation, with comparable linear–quadratic model fits (extra sum-of-squares F-test, p = 0.0923), confirming preservation of the intrinsic radiobiological response under simultaneous irradiation.

**Conclusions:** These results support the reliability of the proposed multidose irradiation system for *in vitro* radiobiology. This robust and reproducible system enables efficient characterization of radiobiological parameters.

## Introduction

*In vitro* radiobiology research plays a vital role in the study of the biological effects of ionizing radiation on living tissues, including the cellular and molecular mechanisms involved in the radiation response, and the radiosensitivity of both normal and tumor cells (Hall and Giaccia 2012; McBride and Schaue 2020). The experimental characterization of dose–response relationships *in vitro* often relies on irradiation protocols that are resource-intensive, time-consuming, and prone to inter-experimental variability, particularly when multiple dose levels are required (Jensen et al. 1989; Nuryadi et al. 2018; Matsui et al. 2019; Brix et al. 2020).

Previous studies have proposed different solutions to this issue. Abatzoglou et al. (2013) proposed a multidose irradiation protocol for 96-well multiwell plates using a clinical linear accelerator; however, the small size of the wells in this plate does not allow the performance of a clonogenic assay, which remains the gold standard assay for evaluating radiation-induced DNA damage. Elliott (2018) developed a high-throughput irradiation device for clonogenic assays, which allows different doses to be delivered within the same 96-well plate with high precision; however, this device is not commercially available nor commonly accessible in radiotherapy research centers, and therefore its protocol would not be easily applicable in most laboratories. Other studies, such as those by Dong et al. (2018) and Mahdavi et al. (2019), also propose irradiation protocols in multiwell plates, but are limited to uniform whole-plate irradiations, in which the entire plate is exposed to a single dose per irradiation session. Additionally, as highlighted by Claridge Mackonis et al. (2018), standard clinical beam qualities and vessel geometries inherently introduce complex dose perturbations that require careful evaluation.

To address these limitations, we propose the development of a robust and reproducible irradiation protocol for multiwell cell culture plates that allows multiple dose levels to be delivered within the same plate, optimizing experimental resources and enabling reliable radiobiological characterization.

## Materials and Methods

### Experimental setup

24–well polystyrene multiwell plates with flat bottoms, sterile (Corning Costar) were used. For both CT imaging and irradiation, solid water slabs (98% polystyrene + 2% TiO_2_, ρ = 1.045 g/cm^3^) (IBA Dosimetry) were placed surrounding the multiwell plate. An equivalent thickness of 5 cm was positioned both below and above the plate to ensure adequate backscatter and build-up regions. Auxiliary elements such as positioning blocks were used to keep the solid water slabs stable and correctly aligned. Figure 1A shows the experimental setup and the position of the multiwell plate. Non-irradiated control samples were processed in parallel, seeded in a separate 24-well plate to avoid unintended radiation exposure. During protocol optimization, some plate configurations included 1% agar or sterile water filling of wells.

**Figure 1.**
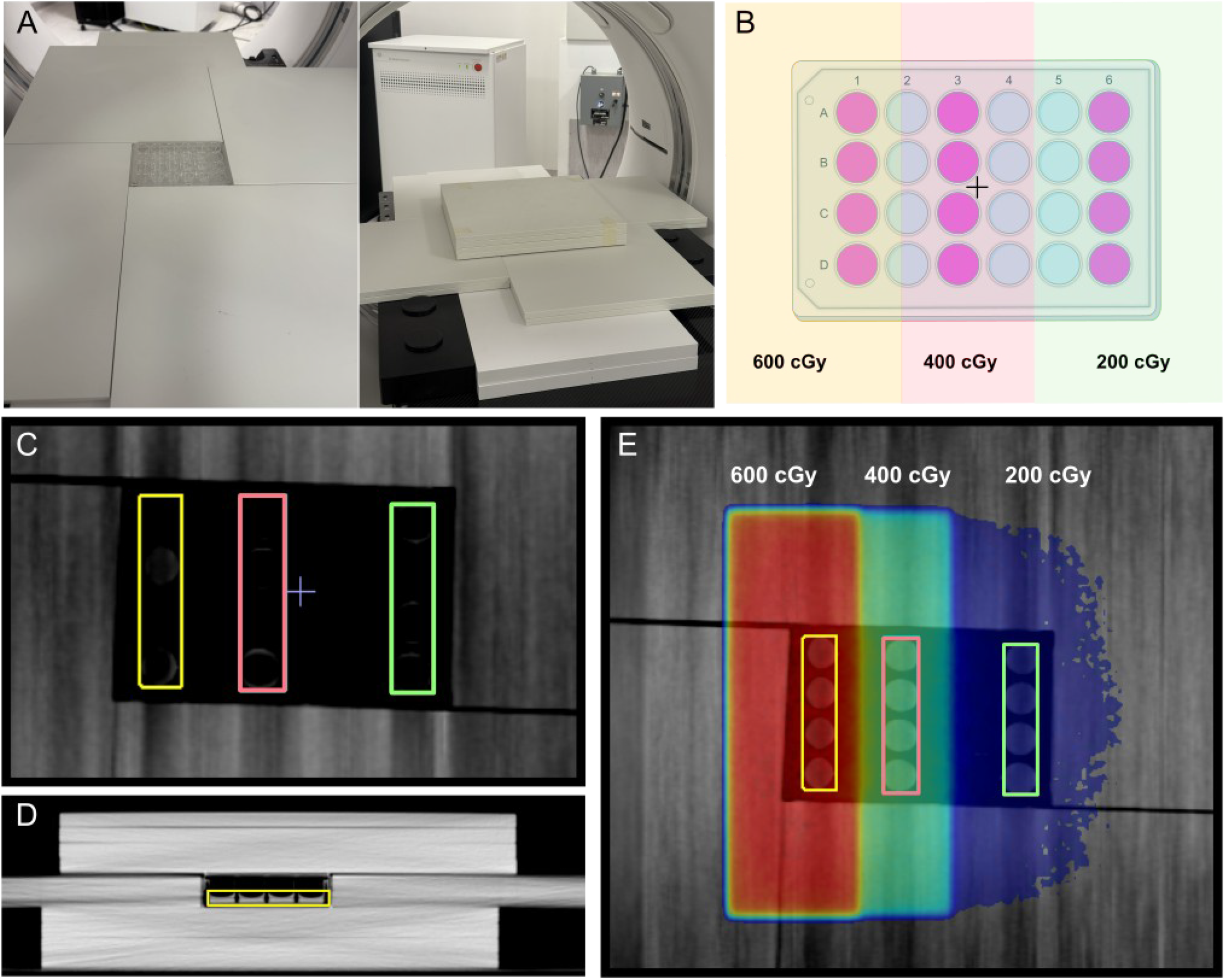
Experimental setup and irradiation planning. **A**. Positioning of the experimental setup for planning CT and irradiation. Placement of the multiwell plate (*left*) and complete setup (*right*). **B**. Layout of the 24-well cell culture plate and irradiation fields. Columns seeded with cells are shown in pink. Each column was assigned to one of three dose levels (200, 400, or 600 cGy) according to the protocol. The cross indicates the plate isocenter. **C-D**. Coronal **(C)** and sagittal **(D)** CT slices depicting rectangular contouring of the entire column at half the height of the wells. **E**. Coronal CT slice showing the three irradiation dose levels (200, 400, and 600 cGy) and the contoured volumes of interest (VOIs).

### Irradiation planning and system

CT images for irradiation planning were acquired using a Discovery CT590 RT scanner (General Electric). Treatment planning was performed using Monaco software version 5.11 at the Radiotherapy Service of INTECNUS Foundation. A 3D-CRT plan was generated using 6 MV X-rays, applying a source-to-axis distance (SAD) technique, with the isocenter defined at the center of the multiwell plate. Dose calculation parameters were set as follows: grid spacing of 0.2 cm, dose deposition to medium, and a statistical uncertainty of 1% per calculation. Irradiations were performed using a clinical linear accelerator (Elekta Synergy Platform).

### Dosimetric verification

To verify the dose calculation, thermoluminescent radiation detectors TLD-100 (Thermo Fisher Scientific), made of lithium fluoride (LiF:Mg, Ti, natural lithium) were used during the irradiation. Custom holders were fabricated via 3D printing to accommodate two detectors and to fit within the plate wells. Volumes of 1% agar were molded to cover the holders, ensuring electronic equilibrium conditions. Crystal readout was performed using a Harshaw TLD 3500 Manual Reader (Thermo Fisher Scientific). In addition, dose verification was carried out using EBT3 radiochromic film (Gafchromic). The film was scanned 20 h after irradiation using an Expression 11000XL scanner (Epson) and processed with myQA software (IBA Dosimetry).

### Biological assessment

#### Cell culture

A breast carcinoma cell line, T-47D, was used. Cells were cultured with RPMI-1640 medium supplemented with 1% penicillin–streptomycin (10,000 U/mL) and 10% fetal bovine serum (FBS), at 37°C and 5% CO_2_.

#### Clonogenic assay

Cells were seeded in 24-well plates at dose-dependent densities (control, 100; 200 cGy, 200; 400 cGy, 400 and 600cGy, 600 cells) and incubated for 24 h before irradiation. After irradiation, cells were cultured at 37°C and 5% CO_2_ for 7 days. Colonies were then fixed using Carnoy’s solution for 15 min, stained with 0.5% w/v crystal violet for 15 min, washed with PBS, and air-dried. Colonies containing more than 50 cells were scored using FIJI software (ImageJ v1.53). Plating efficiency (PE) and surviving fraction (SF) were calculated according to Franken et al. (2006), as follows:

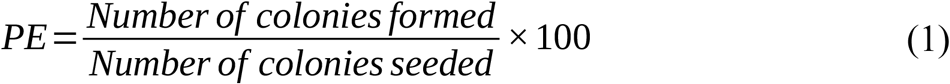

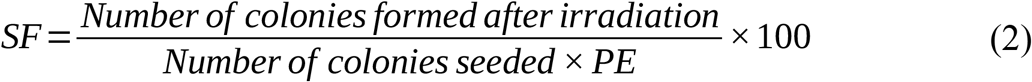

Survival data after irradiation were fitted using a linear–quadratic model (LQM) according to the following equation:

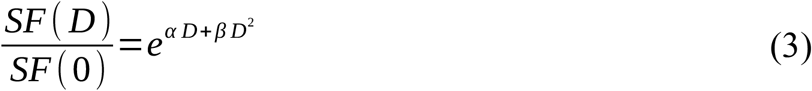

where SF(D) is the surviving fraction at a given dose D, SF(0) is the surviving fraction of the control, and α and β are cell-specific parameters.

#### MTT assay

100 cells per dose level of the T-47D cell line were seeded in a 24-well multiwell plate and irradiated with 200, 400 and 600 cGy, including a non-irradiated control. Cells were incubated at 37 °C and 5% CO_2_ for 7 days. The culture medium was then removed, and cells were washed with PBS. Fresh medium without FBS was then added, together with 50 µL of MTT solution (5 mg/mL in distilled water; M6494, Invitrogen). Plates were incubated for 2 h 30 min at 37 °C and 5% CO_2_. The supernatant was completely removed, and 300 µL of DMSO was added. Absorbance was measured at 540 nm, with background subtraction at 690 nm.

#### γH2AX foci assay

DNA double-strand breaks were detected using the γH2AX foci assay (Rogakou et al. 1998). For this purpose, cells were seeded onto sterile round glass coverslips (12 mm diameter) placed inside the wells of 24-well plates before irradiation. 1:30 h after irradiation, cells were washed and fixed in 4% PFA for 15 min. Cells were blocked with 5% FBS + 0.3% TritonX-100 in PBS for 1 h and incubated overnight at 4°C with primary antibodies (rabbit polyclonal anti-Phospho-Histone H2A.X (Ser139) 1:800 (2577, Cell Signal)), and then with Alexa Fluor 488 (4412, Cell Signal) secondary antibody (1:1000) for 2 h at RT. Hoechst-33258 was used for nuclear staining. Samples were mounted in glycerol, and images acquired on a Confocal Fluorescence Microscope (LSM 980, Carl Zeiss) and processed with FIJI software (ImageJ v1.53).

### Statistical Analysis

Data were analyzed using GraphPad Prism version 8.0.1 (GraphPad Software, CA, USA). Normality was assessed using the Shapiro–Wilk test, and homoscedasticity was evaluated using Bartlett’s test. Cell proliferation data were analyzed by one-way ANOVA followed by multiple comparisons testing. Survival data were fitted using the linear–quadratic model (LQM), and curve comparisons between irradiation conditions were performed using an extra sum-of-squares F-test. Statistical significance was assumed when p < 0.05.

## Results

### Experimental irradiation setup and dose planning

#### Definition of dose volumes

To achieve a graded irradiation, three dose levels were delivered within a single plate by irradiating selected columns separated by empty columns, as shown in Figure 1B. This arrangement provides quadruplicate replicates for each dose level, while avoiding overlap between the irradiated volume and the beam penumbra region.

Dose estimates were obtained under four contouring scenarios: (1) volume of the entire column; (2) volume of the entire column at half the height of the wells, corresponding to the height of the culture medium volume; (3) rectangular contouring around each individual well within a column; (4) manual freehand contouring of the culture medium volume (Supp. Fig. 1A). Each scenario yielded distinct dose values under a uniform 200 cGy open-field irradiation, as reported in Supplementary Table 1. The uniformity of the calculated dose is only minimally affected by the choice of structure selection, with a maximum deviation of 1.81 cGy (0.88%) from the estimated mean dose. Therefore, contouring configuration 2, corresponding to mid-height columns, was selected, establishing a compromise between contouring simplicity and reduced inhomogeneities within the volume of interest (Figure 1C-D).

The impact of multiwell plate filling on dose distribution was also evaluated during the system design phase using both agar and water (Supp. Fig. 2A). Dose inhomogeneity decreased as fillers were added to the plate, indicating that the presence of air within the setup reduces dose uniformity. Mean dose percentage deviations relative to the prescribed values were calculated for each dose level across all planning configurations and remained below 5% in all cases (Supp. Fig. 2B), a range considered unlikely to produce biologically meaningful differences given the intrinsic variability of clonogenic assays. Such variability arises from fluctuations in PE, uncertainties in cell seeding and cell counting, colony scoring procedures, and cell–cell interaction effects, all of which are known to influence survival estimates (Brix et al. 2021). Since the differences between configurations were therefore not expected to produce biologically meaningful effects, and given that plate filling increases the likelihood of handling errors and potential sample contamination, the experimental setup was optimized to improve the performance of the selected configuration, as detailed in the following sections.

#### Field definition and irradiation planning

Two different cell culture configurations were evaluated to achieve the simultaneous delivery of 200, 400 and 600 cGy: a *single-spacing* arrangement, in which one empty column was left between every dose level, and a *double-spacing* arrangement, in which two empty columns were introduced between the 200 and 400 cGy dose regions (Supp. Fig. 3A).

Three irradiation field sizes were defined in the isocentric plane, one for each dose level. Irradiation was delivered such that beam superposition resulted in the prescribed dose for each column, while ensuring dose uniformity within the volume of interest. The same fields were planned with the linear accelerator gantry positioned at 0° and 180°, implementing an anterior– posterior/posterior–anterior (AP–PA) irradiation technique. The 200 cGy column was irradiated only with an open 20 cm × 20 cm field; for the 400 cGy column, an additional 20 cm × 11 cm field was added; finally, for the 600 cGy column, an additional 20 cm × 6.5 cm field was applied.

Dose distributions obtained with the *single-spacing* configuration showed partial overlap between adjacent regions, resulting in increased dose gradients at the interfaces between adjacent regions (Supp. Fig. 3B). In contrast, the *double-spacing* configuration provided improved spatial separation between dose regions, with reduced cross-dose contamination and more homogeneous dose distributions within each region. Based on these results, the *double-spacing* configuration was selected for the final irradiation protocol (Figure 1E). A detailed dosimetric comparison between both configurations is provided in the Supplementary Material. The planning of the multidose irradiation protocol was performed using the defined setup. Resulting gantry angle parameters, field sizes, and monitor units (MU) for each field are detailed in Supplementary Table 2. Based on the optimization process, a final irradiation plan was generated using the double-spacing configuration, delivering three distinct dose levels (200, 400, and 600 cGy) within a single multiwell plate. The resulting dosimetric parameters showed well-defined and spatially separated irradiation regions (Table 1), with dose homogeneity compatible with *in vitro* radiobiology experiments.

**Table 1.** Estimated parameters of the irradiation planning. The minimum, maximum, and mean dose values (D_min_, D_max_, D_mean_) in cGy are listed, together with the volume receiving at least 95% (V95) and 100% (V100) of the prescribed dose, the upper and lower dose ranges expressed as percentages, and the deviation of the mean dose from the stipulated dose value.

| | $D_{\min}(\text{cGy})$ | $D_{\max}(\text{cGy})$ | $D_{\text{mean}}(\text{cGy})$ | $V_{95}(\%)$ | $V_{100}(\%)$ | $+\Delta(\%)$ | $-\square(\%)$ | $\square_{\text{mean}}(\%)$ |
| --- | --- | --- | --- | --- | --- | --- | --- | --- |
| 600 cGy | 580.30 | 632.00 | 609.80 | 100.00 | 86.05 | 3.64 | 4.84 | 1.63 |
| 400 cGy | 379.30 | 426.50 | 404.30 | 99.00 | 68.62 | 5.49 | 6.18 | 1.08 |
| 200 cGy | 190.00 | 210.80 | 200.70 | 100.00 | 67.60 | 5.03 | 5.53 | 0.35 |

### Dosimetric verification

Dosimetric verification was performed using two different approaches: TLDs and radiochromic film (RF). For the first approach, two TLDs were placed per dose level, one in each extreme well of the columns corresponding to each dose, where the highest dose variability was expected. Crystal readout was performed and the measured values were compared with those obtained from the treatment planning software. In all cases, the uncertainty is reported for k = 2 and includes both the calibration factor uncertainty (6%) and the dispersion of each irradiation point along the calibration curve. TLD measurements showed mean dose deviations below 1% for all dose levels, while dose ranges reached up to 22.1% at 200 cGy, reflecting higher spatial variability at lower doses. Measurements for each irradiated dose level are presented in Table 2.

**Table 2.**
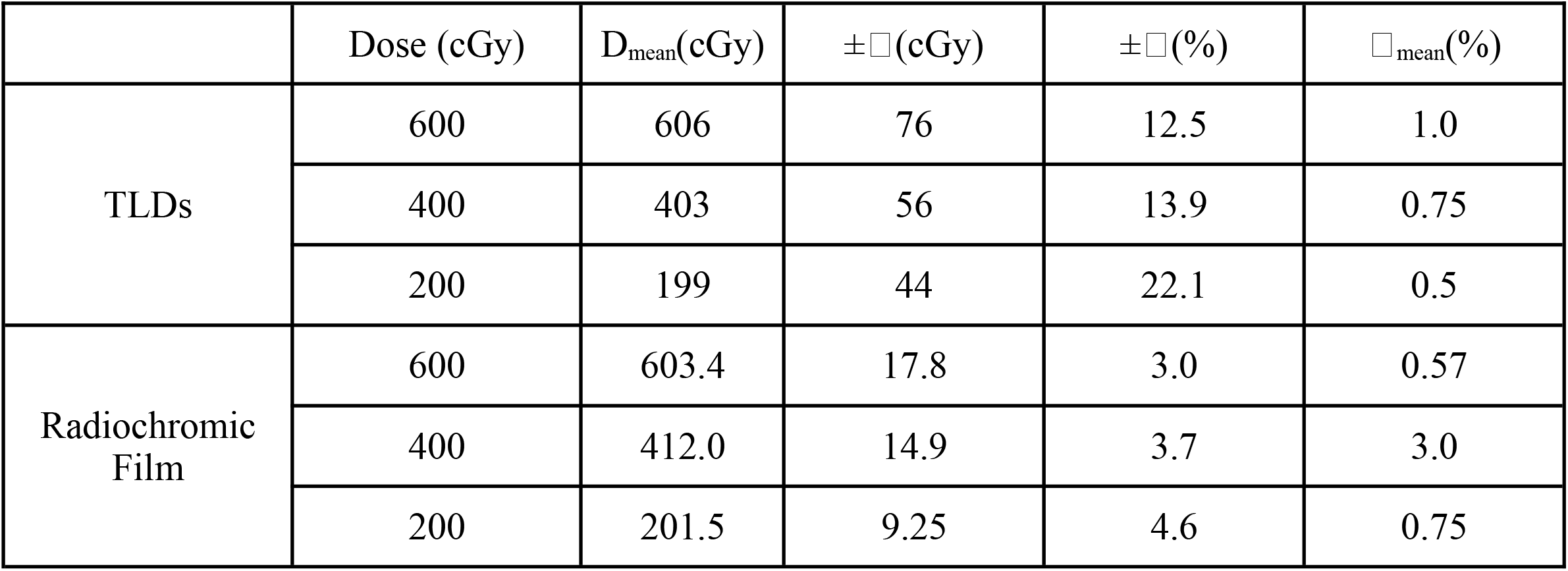
Results of the dosimetric validation of the protocol using TLDs and radiochromic film. The mean dose value (D_mean_) for each theoretical dose level is reported, together with the minimum and maximum variations (±□) in cGy and as a percentage of the prescribed dose, as well as the deviation of the mean (□_mean_) from the stipulated dose value expressed as a percentage.

| | Dose (cGy) | $D_{\text{mean}}$(cGy) | $\pm \square$(cGy) | $\pm \square$(%) | $\square_{\text{mean}}$(%) |
| --- | --- | --- | --- | --- | --- |
| TLDs | 600 | 606 | 76 | 12.5 | 1.0 |
|  | 400 | 403 | 56 | 13.9 | 0.75 |
|  | 200 | 199 | 44 | 22.1 | 0.5 |
| Radiochromic Film | 600 | 603.4 | 17.8 | 3.0 | 0.57 |
|  | 400 | 412.0 | 14.9 | 3.7 | 3.0 |
|  | 200 | 201.5 | 9.25 | 4.6 | 0.75 |

To provide a higher spatial resolution assessment of dose distribution, an RF was placed immediately beneath the multiwell plate, which contained agar volumes simulating the cell culture within the wells. Figure 2A presents a comparison between the dose profile obtained from the treatment planning system (TPS) and that measured using the RF. Measurements showed lower dose dispersion compared to TLDs, with maximum dose ranges of 4.6%, 3.7%, and 3.0% for 200, 400, and 600 cGy, respectively. Mean dose deviations remained below 3% for all dose levels, with the largest deviation observed at 400 cGy (Table 2). Figure 2B shows the comparison between the measured doses and the planned values for both TLDs and RF. Together, these results confirm the accuracy and reproducibility of the dose distribution achieved with the proposed configuration.

**Figure 2.**
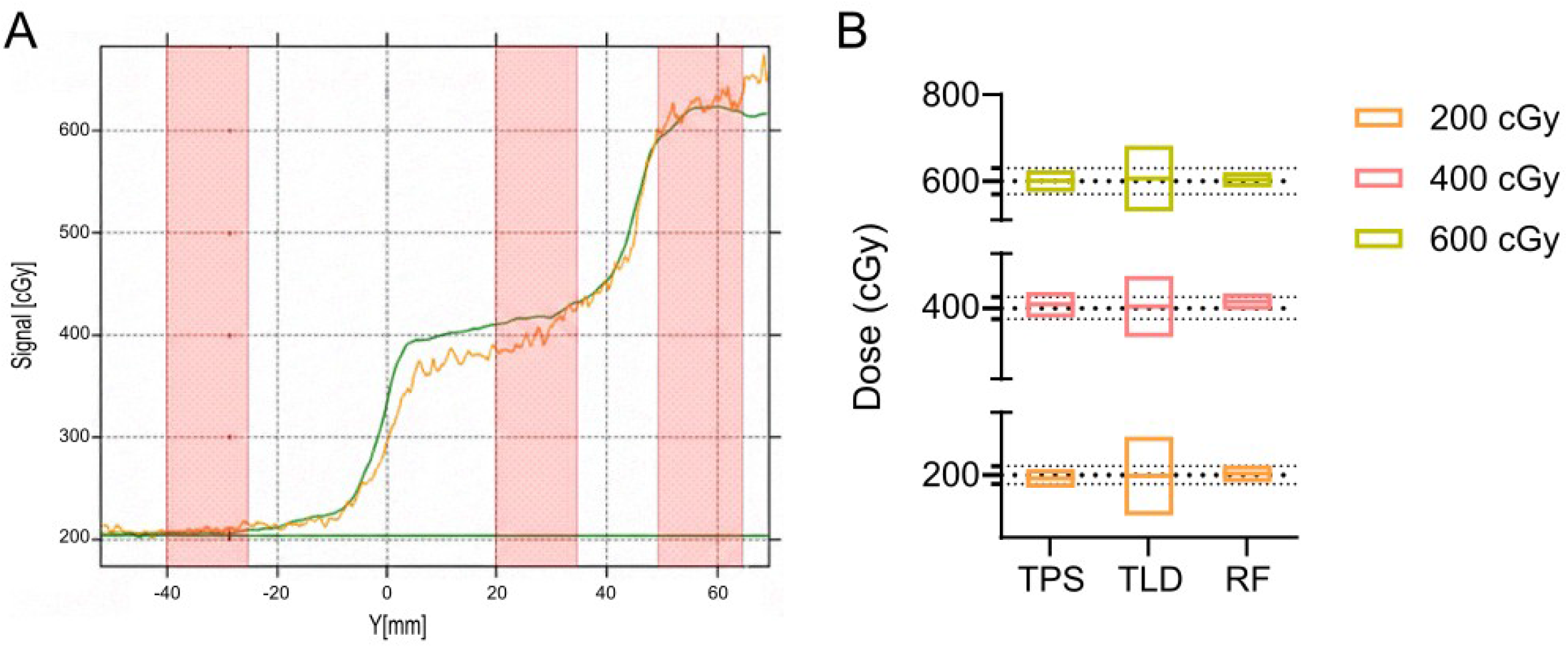
Physical dosimetric verification. **A**. Dose profiles as a function of position along the Y-axis. The green curve represents the values estimated by the Monaco treatment planning system, while the orange curve corresponds to the values measured using radiochromic film (RF). From left to right, the pink bands indicate the estimated positions of the wells receiving 200, 400, and 600 cGy, respectively. **B**. Comparison of the mean, minimum, and maximum dose values obtained from the treatment planning system (TPS) and from dosimetric validation using TLDs and RF.

### Biological assessment

Clonogenic assays were performed under both the multidose and conventional condition, i.e., irradiating a single dose per plate using an open field. A high level of agreement was observed in the fits of the LQM model for both conditions, with SF2 values of 0.7273 and 0.6762, respectively (Figure 3A-B). Comparison of the fitted curves using an extra sum-of-squares F-test indicated that a single shared curve adequately described both datasets (F _(2,23)_ = 2.647, p = 0.0923).

**Figure 3.**
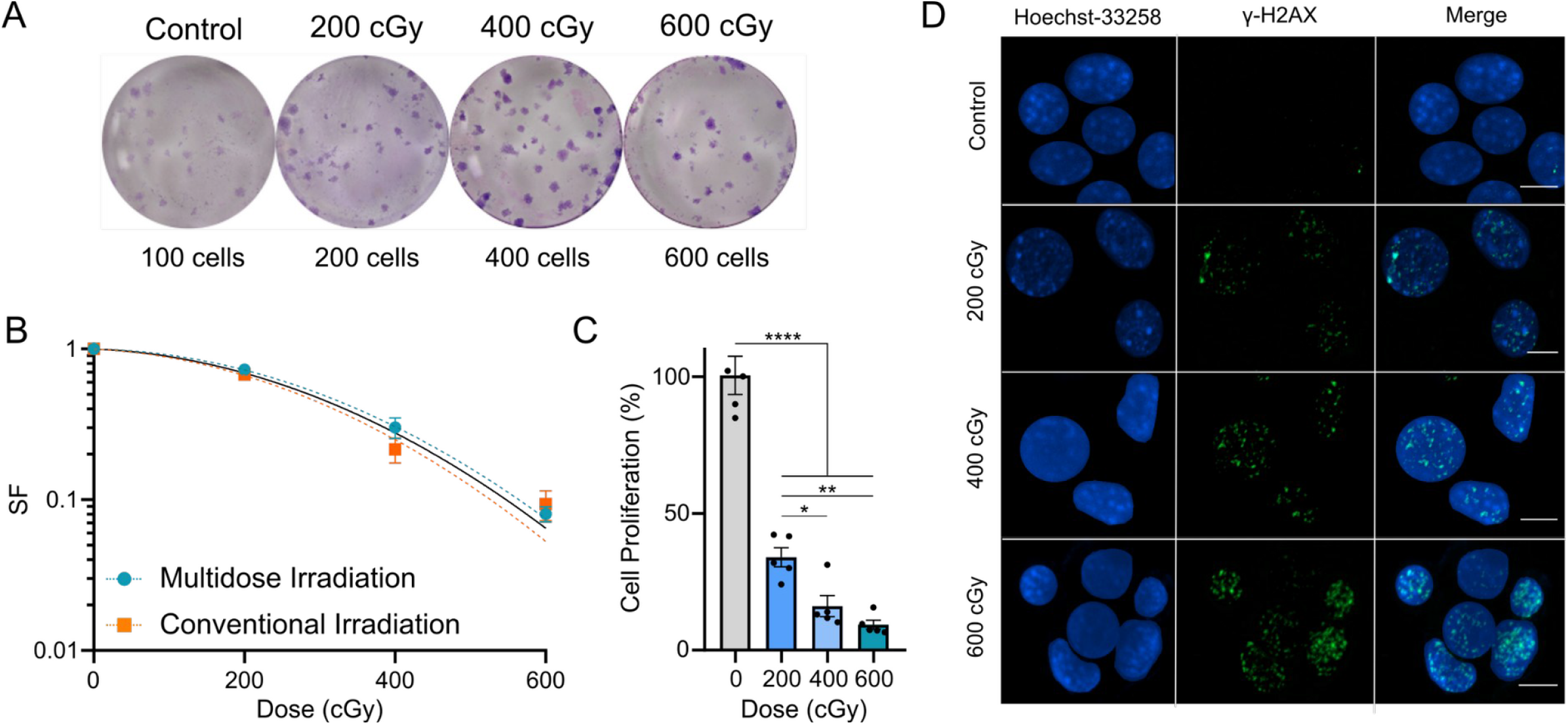
Biological assessment of the multidose irradiation protocol. **A**. Clonogenic assay performed in 24-well plates, with colonies formed by T-47D cells. **B**. Survival curves obtained using the multidose irradiation protocol vs. conventional irradiation. Curves were generated after 7 days post-irradiation for T-47D cells. Each data point represents the mean ± SEM of independent replicates (multidose, n = 3; conventional, n = 4). Dashed lines represent the linear–quadratic model (LQM) fit for each irradiation condition, whereas solid black line represents the shared fit describing both datasets (extra sum-of-squares F-test: F_(2,23)_ = 2.647, p = 0.0923, n.s.). **C**. Cell proliferation of T-47D cells measured by MTT assay 7 days post-irradiation. A significant dose-dependent reduction in cell proliferation was observed following exposure to 200, 400, and 600 cGy compared to the non-irradiated control (one-way ANOVA, F_(3,16)_ = 88.77, p < 0.0001. Holm-Sidak’s multiple comparisons test, **** depicts p < 0.0001, **p < 0.01, *p < 0.05). Data are presented as mean ± SEM (n = 5). **D**. Representative image of γ-H2AX foci (green) and nuclear counterstain with Hoechst-33258 (blue) after irradiation with 200, 400 and 600 cGy. Scale bar, 10 μm.

Consistent with the clonogenic assay results, a significant dose-dependent reduction in cell proliferation was observed following multidose irradiation, in agreement with the delivered dose levels (one-way ANOVA, F_(3,16)_ = 88.77, p < 0.0001) (Figure 3C). Cell proliferation decreased progressively with increasing dose, further supporting the biological effectiveness and internal consistency of the proposed multidose irradiation protocol. Radiation-induced

DNA double-strand damage was qualitatively assessed by immunodetection of γ-H2AX foci following multidose irradiation (Figure 3D). An increase in both the number and intensity of γ-H2AX foci was observed with increasing dose, consistent with the expected dose-dependent induction of DNA damage.

## Discussion

We developed a multidose irradiation protocol for multiwell plates using clinical radiotherapy equipment. The planning yielded good results in terms of dose variability within each dose level, with a maximum variability of 6.18% for 400 cGy and a maximum mean dose deviation of 1.63% for 600 cGy. These results are consistent with the decision to prioritize the 200 cGy dose region, since it corresponds to the standard fraction dose in conventional radiotherapy and therefore represents a clinically relevant radiobiological parameter, achieving a dose variability of 5.53% and a mean dose deviation of only 0.35% for this level. Although the results obtained with the final selected configuration are satisfactory for the purpose of this work, the data show that a plate configuration with filling could be chosen if even greater dose uniformity is desired, as shown in Supp. Fig. 2.

Physical dosimetric verification using thermoluminescent dosimeters and radiochromic films demonstrated high agreement between planned and measured doses across all dose levels. Mean dose deviations remained below 3%, consistent with accepted tolerances for *in vitro* radiobiology experiments. The larger uncertainty observed in TLD measurements is attributable to intrinsic factors of the technique, including calibration factor propagation, limited detector sampling within each dose region, and point-based dose readout in a highly heterogeneous geometry. In contrast, radiochromic films provided planar dose information with reduced dispersion. Nonetheless, both methods confirmed the reliability of the treatment planning–based dose estimation, providing complementary validation of the irradiation protocol.

Biological assessment further confirmed the robustness of the system. Clonogenic assays performed under multidose and conventional single-dose conditions yielded comparable linear–quadratic model fits, demonstrating that simultaneous irradiation does not alter the fundamental radiobiological response. These findings represent a critical validation step, as clonogenic survival remains the gold standard endpoint in radiobiology. Consistent results obtained from both the metabolic assay and DNA double-strand break analysis reinforced these observations, showing a clear dose-dependent reduction in cell proliferation aligned with the delivered dose levels. When comparing the results of this protocol with those reported in the literature, it is observed that previous studies present limitations in terms of dosimetric or biological validation of the developed protocols. Mahdavi et al. (2019) tested different plate filling configurations in 6- and 96-well plates. For this purpose, they used paraffin at the base - requiring an additional sterilization step once the paraffin had dried to allow subsequent cell culture - and saline solution in the internal cavities. They reported differences of up to 37% in the delivered dose between unfilled and filled plates; however, this result was not dosimetrically verified and was only assessed through clonogenic and MTT assays. Abatzoglou et al. (2013) developed a multidose irradiation protocol for 96-well plates using a clinical linear accelerator (LINAC). In this protocol, 8 of the 12 columns of wells were irradiated with graded doses calculated through a system of equations based on dose proportions obtained from dosimetry using TLDs. Biological validation was performed using AlamarBlue metabolic assays, similar to MTT; given the very small well size of this plate format, a graded irradiation protocol based on a 96-well plate is not the most suitable option when the aim is to validate radiation-induced biological effects, such as those assessed by clonogenic assays. Elliott (2018) developed a multidose irradiation protocol using 384-well plates and radiochromic film dosimetry with kilovoltage photon beams (10–225 kVp). Although graded dose delivery was achieved, the reported mean dose deviations reached up to 80 cGy. This limitation is inherently linked to the physical challenges of collimating low-energy kilovoltage beams into millimeter-sized targets, as well as the need for a highly specialized, non-commercial robotic platform to execute the automated positioning and in situ scanning. In contrast, the protocol presented here employs megavoltage photon beams (6 MV) and increased spacing between irradiated columns, resulting in substantially improved dose accuracy and reproducibility, with mean dose deviations below 12 cGy, while maintaining compatibility with clonogenic and other radiobiological assays.

Taken together, the dosimetric and biological results demonstrate that the proposed protocol can accurately deliver multiple dose levels within a single irradiation procedure. Its straightforward implementation further supports reproducible dose-response studies under controlled experimental conditions and facilitates its implementation in other radiotherapy centers.

## Conclusions

The proposed protocol overcomes key limitations of conventional approaches, including high resource consumption, extended operating times, and inter-experimental variability associated with separate irradiations. Physical verification using thermoluminescent dosimeters and radiochromic film demonstrated excellent agreement between calculated and measured doses. Biological assessment through clonogenic and a metabolic assay yielded results consistent with those obtained using traditional irradiation protocols. The simplified experimental setup and the ability to deliver three dose levels with replicates in a single irradiation significantly optimizes workflow and clinical equipment usage. Overall, this robust and reproducible system provides a practical platform for basic radiobiology studies and the systematic characterization of dose-response relationships *in vitro*.

## Notes

**Disclosure statement** No potential conflict of interest was reported by the authors.

**Funding** This work was supported by INTECNUS Foundation and the National Atomic Energy Commission (CNEA), Argentina.

### Competing Interest Statement

The authors have declared no competing interest.

