## Supplementary data for "Development of a Multidose Irradiation Protocol for Clonogenic Assays in Cell Culture Plates"

**Supplementary Material**

**Supplementary Table 1.** Estimated parameters of open-field irradiation at 200 cGy using different contouring methods. The minimum, maximum, and mean dose values in cGy are listed ($D_{min,}$ $D_{max,}$ $D_{mean}$), together with the volume receiving 100% of the prescribed dose (V100)), the upper and lower dose ranges (+$\Delta$, -$\Delta$) expressed as percentages, and the percentage shift of the mean dose relative to the theoretical dose value (□$_{mean}$). VOI, volume of interest.

| Configuration | VOI | $D_{min}$(%) | $D_{max}$(%) | $D_{mean}$(%) | $V_{100}$(%) | +□(%) | -□(%) | □$_{mean}$(%) |
|---|---|---|---|---|---|---|---|---|
| 1 | Full-row | 199.60 | 211.7 | 205.00 | 99.99 | 3.27 | 2.63 | 2.50 |
| 2 | Mid-height row | 199.90 | 212.30 | 205.70 | 100.00 | 3.21 | 2.82 | 2.85 |
| 3 | Rectangular well | 196.90 | 211.30 | 203.30 | 97.66 | 3.94 | 3.15 | 1.65 |
| 4 | Freehand well 1 | 201.20 | 212.30 | 206.40 | 100.00 | 2.86 | 2.52 | 3.20 |
| | Freehand well 2 | 200.20 | 210.90 | 206.20 | 100.00 | 2.28 | 2.91 | 3.10 |
| | Freehand well 3 | 200.40 | 210.70 | 206.50 | 100.00 | 2.03 | 2.95 | 3.25 |
| | Freehand well 4 | 201.90 | 210.90 | 206.70 | 100.00 | 2.03 | 2.32 | 3.35 |

**Supplementary Table 2.** Irradiation parameters of the multidose irradiation protocol. AP denotes anteroposterior irradiations, and PA denotes posteroanterior irradiations. The total number of monitor units (MU) delivered to the culture plate is 608.

| Field | Gantry angle | Field size ($cm^2$) | MU |
|---|---|---|---|
| AP1 | 0° | 20 x 20 | 93 |
| PA1 | 180° | 20 x 20 | 93 |

| | | | |
|---|---|---|---|
| AP2 | 0° | 20 x 11 | 106 |
| PA2 | 180° | 20 x 11 | 106 |
| AP3 | 0° | 20 x 6.5 | 105 |
| PA3 | 180° | 20 x 6.5 | 105 |

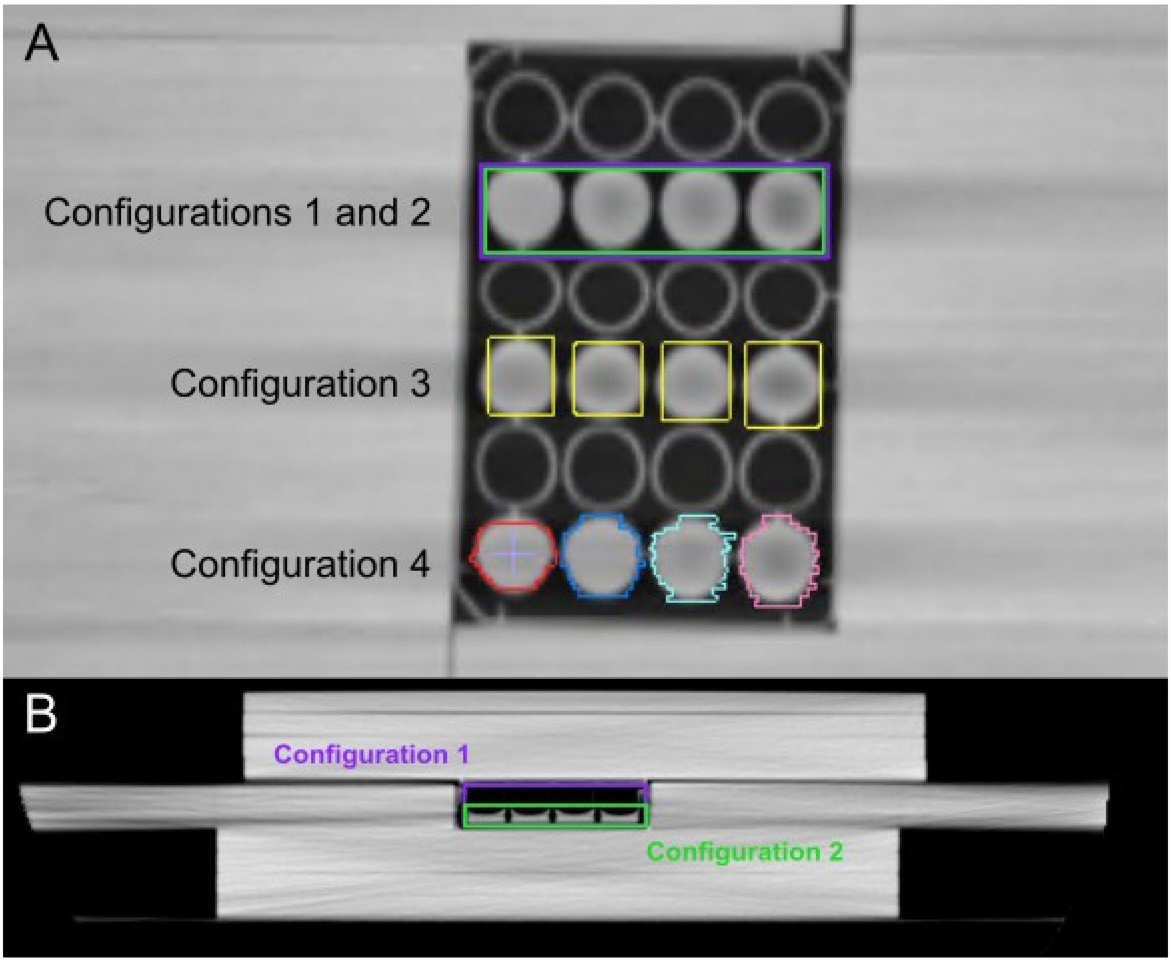


**Supplementary Figure 1. A.** Coronal CT slice showing volume contouring options for dose uniformity evaluation. **B.** Sagittal CT slice showing the height difference between contouring configuration 1 (violet) and 2 (green).

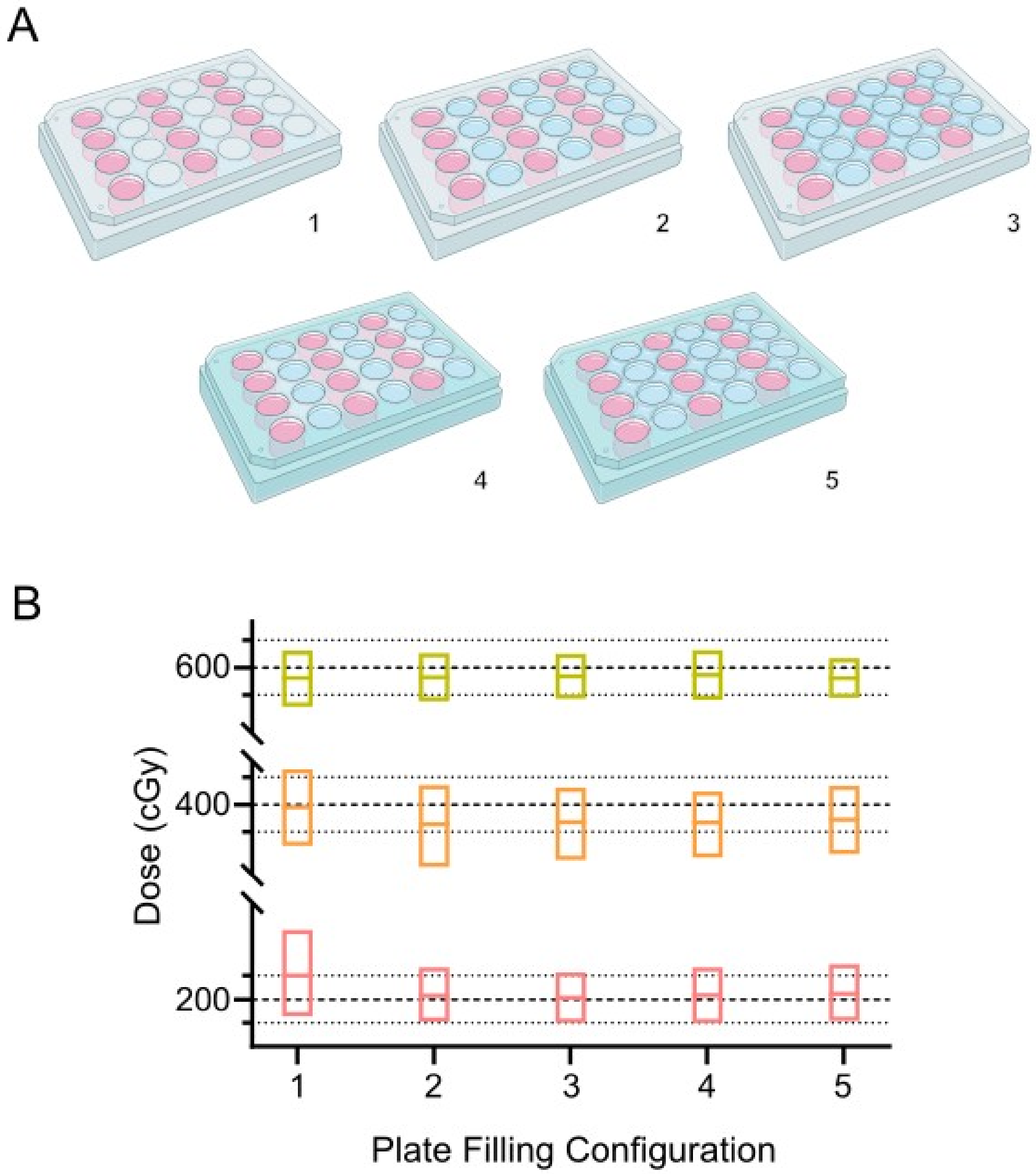


**Supplementary Figure 2. A.** Plate filling configurations: (1) Unfilled plate; (2) plate with water in empty wells; (3) plate with water in empty wells and internal cavities; (4) plate with water in empty wells and agar at the base; (5) plate with water in empty wells and internal cavities, and agar at the base. In all configurations, wells seeded with cells are shown in pink. **B.** Dose variability for each dose range and configuration. Boxes represent the minimum, mean, and maximum dose values obtained from the planning for each dose level. Dashed lines correspond to 200, 400, and 600 cGy, and dotted lines indicate the doses corresponding to 95% and 105% of each dose level.

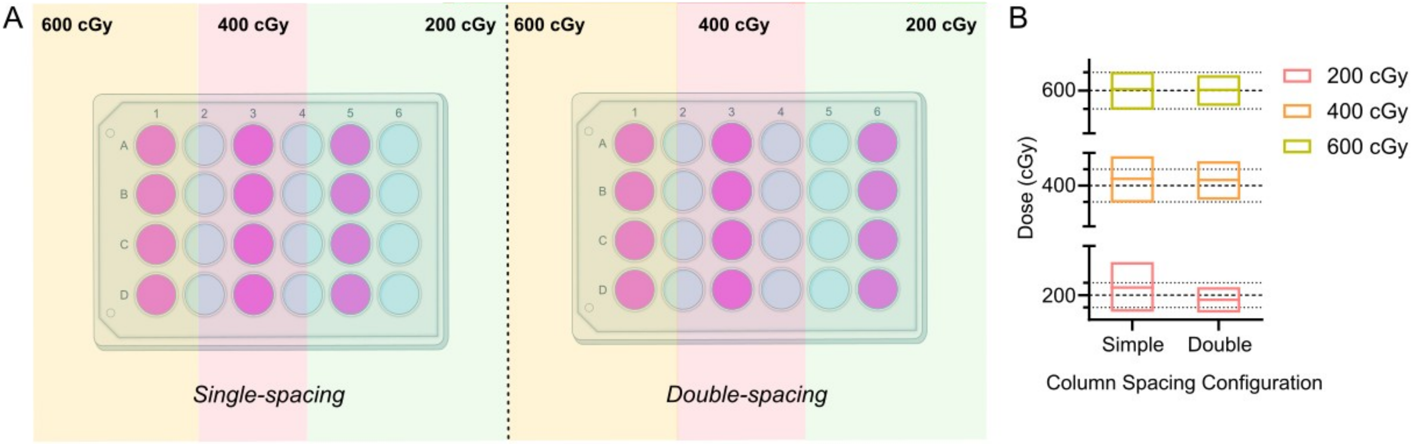


**Supplementary Figure 3. A.** Layout of the 24-well cell culture plate and irradiation fields for multidose irradiation under single-spacing and double-spacing configurations. Columns seeded with cells are shown in pink. Each column was assigned to one of three dose levels (200, 400, or 600 cGy) according to the protocol. **B.** Dose variability comparing single-spacing and double-spacing arrangements. Boxes represent the minimum, mean, and maximum dose values obtained from the treatment planning for each dose level. Dashed lines correspond to 200, 400, and 600 cGy, and dotted lines indicate the 95% and 105% dose levels for each prescribed dose.
